# Cryo-EM structure of detergent-bound *Vibrio cholerae* MakA captures a candidate intermediate between the soluble form and mature assembly of an α-pore-forming toxin

**DOI:** 10.64898/2026.09.08.748681

**Authors:** Yirui Guo, Bradley Quade, Karolina P. Stepien, Tabitha Emde, Alfa Herrera, Raquel Bromberg, Robert Jedrzejczak, Youngchang Kim, Andrzej Joachimiak, Karla J. F. Satchell, Josep Rizo, Dominika Borek, Zbyszek Otwinowski

**Affiliations:** Department of Biophysics, The University of Texas Southwestern Medical Center, Dallas, TX 75390, USA; Ligo Analytics, Dallas, TX 75206, USA; Department of Microbiology-Immunology, Northwestern University Feinberg School of Medicine, Chicago, IL 60611, USA; Center for Structural Biology of Infectious Diseases, Northwestern University Feinberg School of Medicine, Chicago, IL 60611, USA; Structural Biology Center and eBERlight, X-ray Science Division, Argonne National Laboratory, Lemont, IL 60439; Department of Biochemistry and Molecular Biology, University of Chicago, Chicago, IL 60367; Department of Biochemistry, The University of Texas Southwestern Medical Center, Dallas, TX 75390, USA; Department of Pharmacology, University of Texas Southwestern Medical Center, Dallas, TX, USA

## Abstract

The tripartite toxin MakA/MakB/MakE from *Vibrio cholerae* belongs to a family of α-pore-forming toxins that undergo extensive conformational rearrangements upon interacting with membrane. While soluble structures of MakA, MakB and MakE have been determined, the structural basis of membrane association and pore assembly remains poorly understood. Here, we report a cryo-electron microscopy structure of a detergent-associated form of MakA at a global resolution of 3.13 Å, with focused refinement of a flexible region improving the local resolution to 2.68 Å. This structure consists of a trimer of MakA asymmetric homodimers, in which each subunit adopts a transmembrane α-helical conformation distinct from the previously reported soluble structure. The asymmetric homodimer closely resembles the building block observed in the previously reported helical MakA assembly and is structurally similar to the homodimeric building blocks of the *Aeromonas hydrophila* AhlB and *Serratia marcescens* SmhA pores, and the heterodimeric building blocks of the *Xenorhabdus nematophila* XaxAB and *Yersinia enterocolitica* YaxAB pores, suggesting a conserved assembly principle among distantly related α-pore-forming toxins. Complementary hydropathy analysis, AlphaFold2 modeling, and liposome-based assays reveal distinct roles for the three components, with MakA acting as the primary membrane-inserting subunit, MakE exhibiting low but consistent membrane-association, and MakB alone remaining largely soluble. Together, these results support a model in which the MakA asymmetric homodimer represents a plausible early membrane-insertion unit and building block for pore assembly, while the trimer of dimers represents an early assembly intermediate.

## Introduction

The alpha-pore-forming toxins (α-PFTs) are a group of bacterial toxins that are secreted as soluble monomers, and can transform into membrane-bound form and oligomerize when interacting with the eukaryotic cell surface to create pores (1, 2). Such pores are considered virulence factors as they can disrupt eukaryotic cell membrane and lead to ion imbalance and eventually host cell damage (2).

α-PFT can be categorized according to the number of protein components required for pore assembly. A single-component (monopartite) α-PFT consists of a single protein that oligomerizes for pore formation, as in ClyA from *Escherichia coli* (2, 3); a two-component (bipartite) α-PFT is composed of two distinct proteins that assemble into heterodimeric building blocks, as observed in XaxAB from *Xenorhabdus nematophila* and YaxAB from *Yersinia enterocolitica* (4, 5); a three-component (tripartite) toxin utilizes three different proteins to assemble a pore, such as AhlABC from *Aeromonas hydrophila* and SmhABC from *Serratia marcescens* (6, 7). Motility-associated killing (Mak) proteins from *Vibrio cholerae* represent a newly characterized family of cytolytic factors that can assemble into a tripartite toxin (8).

*V. cholerae* employs diverse toxins that promote host colonization and cytotoxicity, many of which disrupt eukaryotic membranes through pore formation (9, 10). Beyond their roles in host damage, these virulence factors may also enhance the environmental fitness of *V. cholerae* by protecting against predation by heterotrophic protozoa (11). These pore-forming toxins often operate as multicomponent systems, requiring the coordinated action of several proteins to assemble into functional membrane-inserted complexes. The Mak toxin proteins (MakA, MakB, and MakE) are encoded on a genomic island within a five gene operon *makDCBAE* (8, 12). The three toxin proteins, MakA, MakB and MakE share modest pair-wise sequence similarity (35-45%; Figure S1) and their soluble forms fold into highly similar structures. The soluble monomers of MakA (PDB: 6EZV, 6DFP) (8, 12), MakB (PDB: 6T8D, 6W1W) (12, 13), and MakE (PDB: 6TAO, 6W08) (12, 13) adopt an α/β-fold architecture that resembles the fold of other α-PFTs in their soluble forms, including AhlB (PDB: 6GRK) (6) and SmhA (PDB: 7A27) (7). The genomic colocalization and structural similarities of MakA, MakB, and MakE collectively support their classification as a tripartite toxin system. MakC (PDB: 8RQY) (14) and MakD (PDB: 6TCT) (14) lack sequence and structural similarity to MakA, MakB, and MakE, localized within the *V. cholerae* cytosol, and are therefore thought to function only in auxiliary or regulatory roles (14).

The current proposed mechanism of MakABE pore assembly is that MakA, MakB, and MakE are secreted as soluble monomeric proteins through the flagellar channel in a proton motive force-dependent manner (8), after which they undergo conformational rearrangements upon interacting with the host cell membrane to assemble into a pore-forming complex (15).

Although high-resolution structures of the soluble forms of MakA, MakB and MakE have been reported in several studies (8, 12, 13), information on their membrane-associated states remains limited. To date, only one structure captures MakA in a non-globular form (PDB: 7P3R) (16). However, in this structure, MakA forms a helical assembly of unclear function, making it difficult to infer mechanistic understanding of how MakA transitions from its soluble form into a pore.

In this study, we determined the structure of detergent-associated MakA, which assembles as a trimer of asymmetric dimers, with trimerization mediated by interactions involving the transmembrane head region of each dimer. This asymmetric dimer closely matches the structural building blocks observed in several distantly related α-PFTs, revealing unexpected architectural similarity despite limited sequence conservation. These findings provide a structural framework for understanding MakA oligomerization and support a model in which the trimer-of-dimers represents an early intermediate in MakABE pore assembly. The structure also identifies interprotomer interfaces that could be targeted by small molecules to destabilize MakA in its detergent-associated state, providing a potential strategy for disrupting membrane insertion and inhibiting the early stages of pore assembly.

## Methods

### Cloning and purification

Cloning, expression and purification of *V. cholerae* MakA, MakB and MakE were described previously (12). The proteins were first purified with an N-terminal 6×His-tag. The His-tag was subsequently removed by cleavage with tobacco etch virus (TEV) protease and Ni-NTA aridity chromatography. The purified tagless protein was dialyzed against buffer containing 20 mM HEPES (pH 7.5), 200 mM NaCl, and 1 mM tris(2-carboxyethyl)phosphine hydrochloride (TCEP). n-dodecyl-β-D-maltoside (DDM) was added to a final concentration of 0.03% to induce the transition to the detergent-associated form. Purified proteins were stored at −80 °C until use.

### Cryo-EM grid preparation and data collection

Quantifoil R 1.2/1.3 300 Mesh Gold grids were glow discharged for 90 s at 30 mA with a PELCO easiGlow™ Glow Discharge Cleaning System to obtain a hydrophilic surface. 3 µl of 18 mg/mL MakA was applied to the glow-discharged surface of the grid at 4 °C, 100% humidity and was blotted for 6 s with blot force 20 using a Thermo Scientific Vitrobot Mark IV System.

The data were acquired with a 300 kV Titan Krios G2 microscope (Thermo Fisher) equipped with a K3 Summit direct electron camera (Gatan) run in super-resolution mode at a nominal magnification of 105,000×, with a physical pixel size of 0.833 Å. A phase plate was not used and the objective aperture was not inserted. SerialEM was used for automated data collection in beam-image shift mode with 9 images collected per stage movement, with a defocus range from −0.8 µm to −2.2 µm and beam-image shift compensation (17). The slit width of the GIF Quantum Energy Filter was set to 25 eV. Movies were dose-fractionated into 125 frames with a total dose of ∼91 e^-^/Å^2^.

### Cryo-EM data processing

All movies were imported into CryoSPARC followed by patch motion correction using binning of 2 and patch CTF correction (18). Particles were first picked using blob picker with diameter range of 200-400 Å on the first 500 micrographs. Picked particles were extracted with box size of 432 pixels Fourier cropped to 216 pixels, and cleaned by several rounds of 2D classifications. Classes showing clear protein features were used as templates for the TOPAZ training (19). 390,971 particles were then extracted using the trained model from all micrographs in 432-pixel boxes Fourier-cropped to 216 pixels.

A total of 304,188 clean particles were retained after one round of 2D classification. These particles were subjected to *ab initio* reconstruction with three classes, followed by heterogeneous refinement, which yielded one class containing 236,459 particles with well-resolved structural features. The selected particles were re-extracted in 432-pixel boxes and refined using non-uniform refinement with C3 symmetry, resulting in a reconstruction at 3.13 Å resolution (global map) (20).

Due to relative flexibility between repeating units, which resulted in blurred density in the distal regions of the map, particle subtraction was performed on symmetry expanded particles to isolate a single repeating segment of the assembly. The resulting particles were subjected to local refinement using a soft mask encompassing this segment, yielding a reconstruction at 2.68 Å resolution. The local and global maps were aligned to a common coordinate system for model building (Table 1).

**Table 1.** Cryo-EM data collection and atomic model building.

|  | **Data collection** |
| --- | --- |
| **Sample name** | MakA |
| **Instrument** | Titan Krios G2 |
| **Detector** | K3 Summit |
| **Energy filter** | BioQuantum |
| **Objective aperture** | No |
| **Nominal magnification** | 105,000× |
| **Data collection mode** | Beam-Image Shift; 3×3 holes, 1 shot/hole |
| **Electron dose rate (e<sup>-</sup>/pixel/s)** | 12.5 |
| **Exposure time (s)** | 5 |
| **Detector pixel size (Å)** | 0.833 |
| **Data pixel size (Å)** | 0.4165 |
| **Movies acquired** | 2,234 |
|  | **Reconstruction** |
| **Molecular weight (kDa)** | 39 × 6 = 234 |
| **Reconstruction symmetry** | C3 |
| **Particles used in reconstruction** | 236,459 |
| **Resolution FSC<sub>0.143</sub> (Å)** | 3.13 (global)/2.68 (local) |
|  | **Atomic model building** |
| **Non-hydrogen atoms** | 15,201 |
| **Protein residues** | 2,052 |
| **Ligands** | 0 |
| **RMSD bond lengths (Å)** | 0.014 |
| **RMSD bond angles (°)** | 1.035 |
| **Model-to-map CC (mask)** | 0.88 |
| **MolProbity score** | 1.00 |
| **Clashscore (all atom)** | 2.25 |
| **Poor rotamers (%)** | 0 |
| **Ramachandran (%)** |  |
| **favored** | 98.48 |
| **allowed** | 1.52 |
| **outliers** | 0 |

### Model building

Initial atomic models were obtained by docking the PDB models of soluble (PDB: 6EZV) and helical form (PDB: 7P3R) of MakA to the maps using MOLREP (21). The distal region was rebuilt in Coot and refined with *Phenix.real_space_refine* and Servalcat against the higher resolution local map (22–29). The resulting local model was then merged with the remainder of the structure and expanded using C3 symmetry to fit the global map. The complete model was further rebuilt in Coot and refined using *Phenix.real_space_refine* and Servalcat. Model statistics were validated using Molprobity server and are summarized in Table 1 (30). Visualization of the structure is done using PyMOL and ChimeraX (31–33).

The atomic coordinate for the model (pdb_000038ke), the composite (EMD-78871), local (EMD-78869) and global (EMD-78868) maps have been deposited in the Protein Data Bank (http://wwpdb.org/) and the Electron Microscopy Data Bank (https://www.ebi.ac.uk/emdb/), respectively.

### Structural analysis

Surface electrostatics analysis was performed using the APBS PyMOL plugin, with default settings corresponding to pH 7, and PDB2PQR was used to add hydrogen atoms and assign protonation states, atomic charges, and radii (34). APBS webserver was used to calculate per residue pKa (34). PDBsum and PISA server were used to analyze binding interface (35, 36). The ProtScale tool from ExPASy was used to calculate the Kyte-Doolittle hydrophobicity (hydropathy) profile of the protein sequence (37, 38). Structural similarity was assessed by calculating the root-mean-square deviation (RMSD) of Cα atoms following structural superposition using PyMOL and Coot (22). PyMOL was used for overall structural superposition, while Coot was used for least-squares (LSQ) superposition and RMSD calculation of selected structural regions.

### AlphaFold modeling

The AlphaFold models were generated using the AlphaFold2 Collab implementation: https://colab.research.google.com/github/sokrypton/ColabFold/blob/main/beta/AlphaFold2_advanced.ipynb with default settings (num_models = 5, ptm option, num_ensemble = 1, max_cycles = 3, num_relax = 0, tol = 0, num_samples = 1) (39). Both chains of the detergent-associated MakA homodimer were used individually or together as templates for modeling the MakB homodimer, MakE homodimer, MakA/MakB heterodimer, MakA/MakE heterodimer and MakB/MakE heterodimer.

### Liposome assays

The epithelial lipid mixture (28% phosphatidylcholine, 27% phosphatidylethanolamine, 0.4% phosphatidylglycerol, 6% phosphatidylserine, 5% phosphatidylinositol, 0.2% phosphatidic acid, 7% sphingomyelin, 25% cholesterol, and 1.4% cardiolipin) was resuspended in the assay buffer containing 20 mM sodium acetate (pH 5.0), 200 mM NaCl, 1 mM TCEP, 10% glycerol, 20% β-octyl glucoside (β-OG) and 50 mM sulforhodamine B (SRB). Liposomes with encapsulated SRB molecules were formed using 100 nm spin columns. To remove excess SRB and isolate properly formed liposomes, co-flotation through a three-layer Histodenz gradient (35%, 25%, and 0%) was performed. Liposomes were harvested from the topmost layer.

For liposome co-sedimentation assays, liposomes (0.5 mM total lipid) were incubated for 2 h at 37 °C with either MakA, MakB or MakE individually (5 μM), or an equimolar mixture of MakA, MakB and MakE (5 μM total protein) that had been preincubated for 2 h at room temperature. Samples were then subjected to ultracentrifugation at 15,000 × g for 60 minutes at 4 °C to separate liposome-bound and unbound protein. The supernatant fraction was collected, and the pellet was washed twice and further purified by two additional rounds of ultracentrifugation under the same conditions. Following the final centrifugation step, the pellet was resuspended in a volume equivalent to that of the collected supernatant. Equal volumes of supernatant and pellet fractions were analyzed by SDS-PAGE.

For liposome co-flotation assays, liposomes (0.5 mM total lipid) were incubated with 10 μM total protein (MakA, MakB, MakE or equimolar mixture of all three proteins) for 2 h at 37 °C. Following incubation, samples were mixed with Histodenz (40%) and loaded beneath a discontinuous Histodenz gradient (40%, 35%, 30%, 0%). Gradients were centrifuged at 280,000 × g for 4 h in an ultracentrifuge. After centrifugation, the top fraction containing floated liposomes was collected and analyzed by SDS-PAGE.

For liposome leakage assays, liposomes containing encapsulated SRB molecules (0.1 mM total lipid) were mixed with 5 μM total protein (MakA, MakB, MakE or equimolar mixture of all three proteins) for 2 h at 37 °C. Fluorescence dequenching of SRB (excitation at 560 nm, emission at 590 nm) was monitored over time at 37 °C. Maximum fluorescence was induced by adding 1% β-OG at t = 2,000 s to release SRB from all liposomes.

### Sequence alignment and evolutionary analysis

Pairwise sequence identity and similarity was calculated using the Needle program from the EMBOSS webserver (40). PSI-BLAST (Position-Specific Iterated BLAST) (41) was used to detect distant homologs and Foldseek (42) was used to search for structurally similar proteins.

A phylogenetic analysis of α-PFTs was performed using a curated set of representative sequences. The dataset included several well-characterized α-PFTs, including proteins with available experimentally determined structures. Homologous sequences were identified through searches of the UniProt database, and 1-2 homologs from deferent species were selected for each reference protein (43). Sequence selection was restricted to homologs sharing 40-80% amino acid identity with the corresponding reference protein to provide broad phylogenetic representation while avoiding excessive sequence redundancy. The sequences were aligned using the MAFFT online server, and phylogenetic analysis was performed using the Neighbor-Joining method based on conserved sites in the alignment (44, 45). Evolutionary distances were calculated using the Jones-Taylor-Thornton (JTT) amino acid substitution model with default settings (rate heterogeneity among sites α = ∞, and no bootstrap resampling). The resulting phylogenetic tree was visualized for comparative analysis of sequence relationships among α-PFTs. Visualization of the multi-sequence alignment was done using Jalview (46).

## Results

### Overall structure of detergent-associated MakA

The detergent-associated MakA structure solved in this study consists of a trimer of asymmetric dimers arranged around a 3-fold symmetry axis (Figure 1a-c, Figure S2). The head regions of the three dimers (chains AB, chains CD and chains EF) are embedded in a compact DDM micelle and interact closely to form the transmembrane core of the assembly. The disk-like DDM density surrounding this transmembrane core is ∼35 Å thick. Because the head regions are inserted at an angle, the length of the helices buried within the detergent is ∼44 Å in chain A (C, E) and ∼38 Å in chain B (D, F) of the asymmetric dimer (Figure 1b).

**Figure 1.**
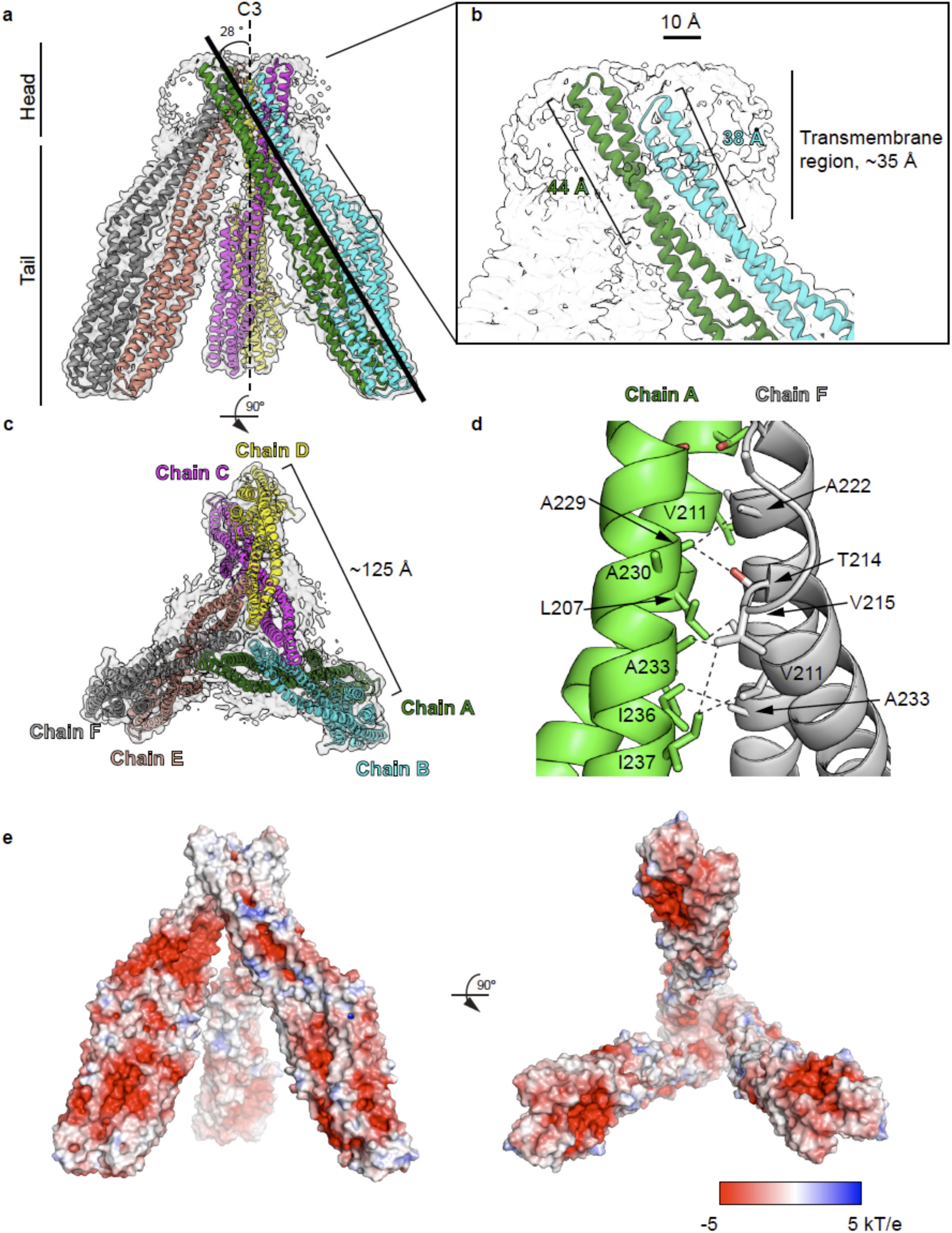
Detergent-associated MakA trimer of dimers. **a)** Cryo-EM map of the MakA trimer of dimers fitted with the atomic model. The head and tail regions are labeled, and the angle between one dimer and the C3 symmetry axis is indicated. **b)** Zoomed-in view of the transmembrane region. Chains A and B, which form the same asymmetric dimer, exhibit slightly different degrees of insertion into the detergent micelle. **c)** Rotated view of the MakA trimer of dimers. **d)** Interaction between two adjacent dimers through chains A and chain F. **e)** Surface electrostatic potential of the MakA trimer of dimers.

The tail region of each dimer is ∼100 Å in length, extending away from the detergent micelle to form a tripod-like assembly. At the distal ends of the legs, the dimers are separated by ∼125 Å (Figure 1c). Each leg is positioned at an angle of approximately 28° relative to the 3-fold symmetry axis (Figure 1a). The tripod is stabilized by interactions at the head region among the asymmetric dimers (chains AB, CD, and EF), involving contacts between chains A-F, B-C and D-E. Each interface (e.g., between chain A residues 204-237 and chain F residues 211-233) consists of a network of hydrophobic interactions, with a buried surface area of 476.6 Å², corresponding to less than 3% of the total surface area of each chain (∼18,000 Å^2^) (Figure 1d). Although each interface is small, every dimer is effectively wedged between the other two, such that the three dimers mutually constrain one another in a closed triangular arrangement. This interlocking geometry restricts relative motion among the head regions of adjacent dimers. In contrast, the distal tail regions showed significant blurring in the cryo-EM density map, likely due to their substantially greater conformational freedom (Figure S2c). Local refinement using symmetry-expanded particles recovered high-resolution features in this region, indicating that the blurring arises from slight differences in the orientation of the tail regions across particles rather than from structural disorder within the subunits themselves (Figure S2d-f).

The observed detergent-associated arrangement of MakA is consistent with the electrostatic surface analysis, with the predominantly hydrophobic transmembrane head region buried within the detergent and the soluble tail region, which contains extensive negatively charged patches, remaining exposed to the aqueous environment (Figure 1e).

In its current configuration, the MakA assembly does not form a pore, as the three dimeric units do not enclose a continuous central channel. However, the open arrangement leaves sufficient space for the addition of further membrane-associated building blocks, which, together with subsequent rotational rearrangements of the dimeric subunits, could give rise to a complete transmembrane pore.

### The asymmetric dimer as a building block

Each asymmetric dimer is composed of two MakA molecules adopting markedly different conformations, both of which direr substantially from the soluble form of MakA in their secondary and tertiary structures (Figure 2a-g). In contrast to the soluble form, in which a β-tongue-like structure folds against a bundle of α-helices, both chain A and chain B of the detergent-associated MakA are composed entirely of α-helices, although the positions of the connecting turns direr between the two chains (Figure 2b). For example, the transmembrane regions adopt deferent conformations in the two chains. In chain A, the two transmembrane helices are connected by a turn comprising residues 217-222, whereas in chain B the corresponding turn spans residues 215-220. This subtle deference leads to distinct inter-helical packing arrangements in the two chains, with the packing within chain A shifted by one or two helical turns relative to that within chain B. (Figure S3). In addition, chain B contains more helix-disrupting breaks and turns, resulting in a greater number of shorter helical segments compared with chain A (Figure 2b-d).

**Figure 2.**
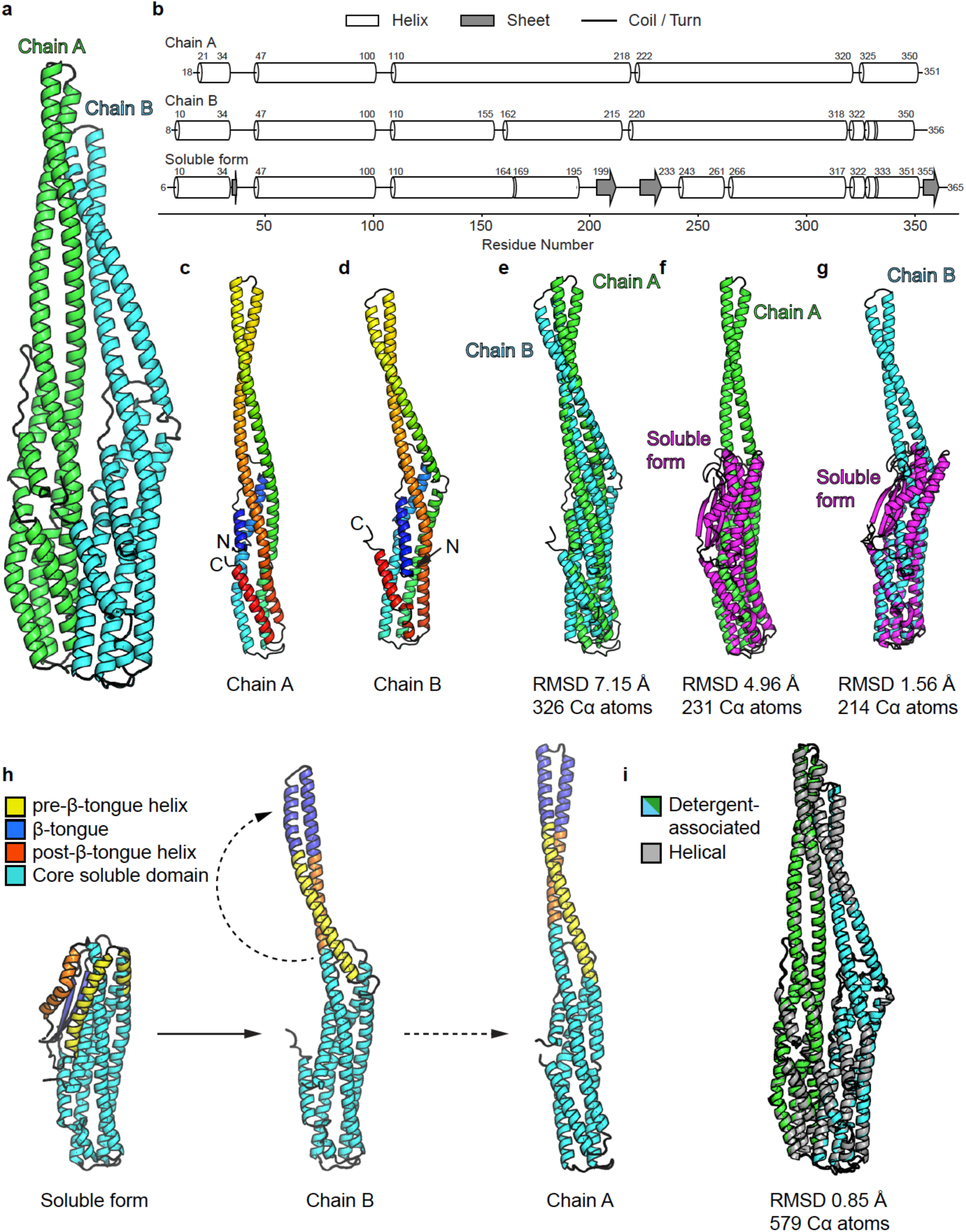
MakA asymmetric dimer. **a)** Atomic model of one asymmetric dimer consisting of chain A (green) and chain B (cyan) from the detergent-associated MakA structure. **b)** Comparison of the secondary structures of chain A, chain B from the detergent-associated MakA structure with the MakA soluble form. **c)** Chain A colored in a rainbow gradient from the N-terminus (blue) to the C-terminus (red). **d)** Chain B colored in a rainbow gradient from the N-terminus (blue) to the C-terminus (red). **e)** Superimposition of chain A and chain B. **f)** Superimposition of chain A with the MakA soluble form (PDB: 6EZV). **g)** Superimposition of chain B with the MakA soluble form (PDB: 6EZV). **h)** Proposed structural rearrangement from the soluble form to chain B and then to chain A. The transition from soluble to transmembrane form of MakA chain B involves the rotation of the pre- and post-β-tongue helices (yellow and orange, respectively) away from the core soluble domain (cyan), as well as the extension and refolding of the β-tongue (blue) into α helices. **i)** Superimposition of the detergent-associated MakA asymmetric dimer with the asymmetric dimer from the MakA helical assembly (PDB: 7P3R).

The superimposition of chain A and chain B resulted in an RMSD of 7.15 Å over 326 Cα atoms, reflecting substantial structural variation (Figure 2e). Superimposition of chain A or chain B onto the soluble form of MakA yielded RMSDs of 4.96 Å over 231 Cα atoms and 1.56 Å over 214 Cα atoms, respectively. Although difference between the β strands in the soluble form and the α helices in chain B (residues 156-265 in both structures) remain significant, the rest of chain B aligns closely with the soluble form (Figure 2f, g). A more controlled superimposition using the same Cα atoms (residues 18-155 and 265-352) of chain A or B against the soluble form yielded RMSDs of 5.26 Å and 2.16 Å, respectively, confirming the greater structural similarity between chain B and the soluble form. The transition from the soluble form to chain B involves only rotation of the β-tongue and its flanking helices (the pre- and post-β-tongue helices), followed by extension and refolding of the β-tongue into α-helices (Figure 2h, Figure S1). The structural similarity between the soluble form and chain B suggests that chain B may represent an early intermediate in the transition toward the detergent-associated state, which may subsequently rearrange into chain A (Figure 2h).

Chain A and chain B bury a surface area of 2,748.8 A^2^, corresponding to 14% of the total surface area of chain A and 16% of the total surface of chain B (36). Interactions between the two chains are predominantly hydrophobic and van der Waal’s interactions, distributed evenly along the interface (Figure S4a-f). Several hydrogen bonds and salt bridges are also present (Figure S4a, g-i).

Unlike many asymmetric dimers that retain an approximate pseudo-2-fold relationship, the two MakA protomers adopt an approximately parallel orientation and form a directional, highly asymmetric interface (Figure 2a). This arrangement suggests that the two conformations are mutually stabilizing, such that adoption of one conformation may favor the complementary state in the neighboring protomer, promoting formation of an asymmetric A-B dimer over a symmetric A-A or B-B dimer. Furthermore, because the A-B interface is inherently asymmetric, it cannot be equivalently propagated to generate an extended “A-B-A-B-…” assembly, supporting the asymmetric A-B dimer as a discrete building block.

This asymmetric dimer closely resembles the building block of the previously reported helical MakA assembly determined at resolution of 3.65 Å, with a global RMSD of 1.94 Å over 668 Cα atoms and a core RMSD of 0.85 Å over 579 Cα atoms, despite the completely deferent higher-order architectures of the two assemblies (trimer-of-dimers vs. helical filament) (Figure 2i) (16). While the overall fold of the building block is highly similar, the higher-resolution structure presented here (3.13 Å global, 2.68 Å local) enables more detailed characterization of intermolecular interactions and conformational features within the dimer, including clearer visualization of side-chain densities. The strong structural similarity at the level of the dimeric building block suggests that the conformation observed in this study is unlikely to be an artifact of detergent stabilization.

### Other α-PFT sharing a MakA-like asymmetric dimer as a structural building block

The asymmetric homodimer of MakA is not unique among homologs (Figure 3). A Fold seek search using the MakA homodimer as a query identified AhlB (PDB: 6GRJ, 6H2F; 40% sequence similarity to MakA with 79.5% coverage) (6) and SmhB (PDB: 7A0G; 33.6% sequence similarity to MakA with 61.7% coverage) (7) among the top hits (E-value ∼10⁻¹¹). Both proteins form pores composed of pentamers of asymmetric homodimers, indicating that the MakA dimer architecture is conserved within this family (Figure 3ab).

**Figure 3.**
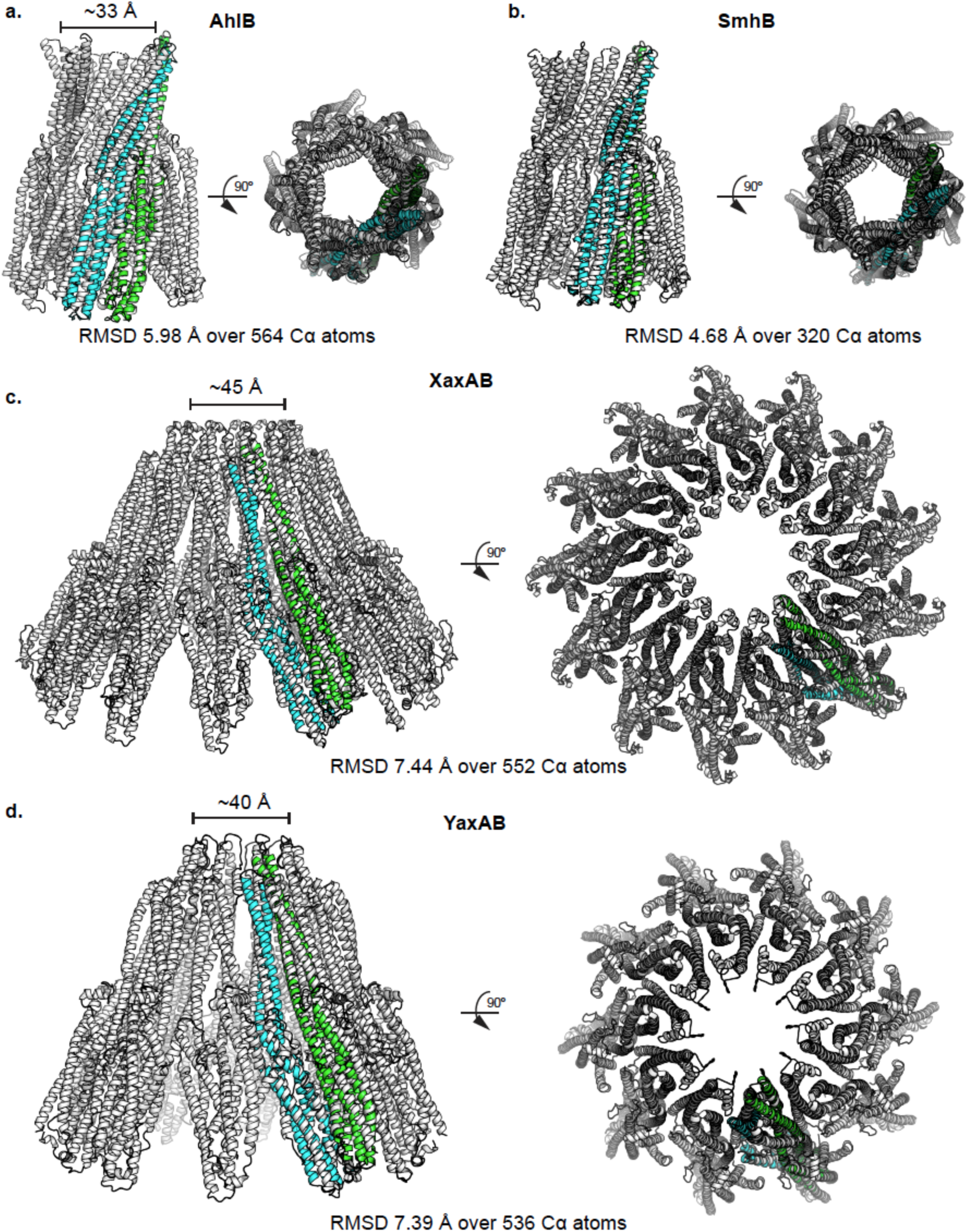
Other α-pore-forming toxins that use asymmetric dimers as building blocks. **a)** Crystal structure of *A. hydrophila* AhlB in a pore-like assembly (PDB: 6GRK). **b)** Crystal structure of *S. marcescens* SmhA (PDB: 7A27) in a pore-like assembly. **c)** Cryo-EM structure of *Y. enterocolitica* YaxAB in a pore-like assembly (PDB: 6EL1). **d)** Cryo-EM structure of *X. nematophila* XaxAB in a pore-like assembly (PDB: 6GY6). In each panel, the MakA asymmetric dimer (chain A: green; chain B: cyan) is superimposed onto the corresponding structure.

More distantly related pore-forming toxins, including XaxAB (PDB: 6GY6) (4) and YaxAB (PDB: 6EL1) (5), were also identified (E-value ∼10⁻⁶). Unlike MakA, AhlB, and SmhB, these toxins assemble as oligomers of heterodimers, with XaxAB composed of 13 XaxA/XaxB heterodimers and YaxAB composed of 10 YaxA/YaxB heterodimers. Although XaxA/XaxB and YaxA/YaxB are not readily detected by conventional sequence-based searches using MakA as a query, they can be identified using iterative profile-based methods such as PSI-BLAST. Visual inspection of the superimposed structures reveals strong conservation of the overall domain architecture between MakA and the XaxA/XaxB or YaxA/YaxB heterodimers despite of RMSD values of 7.44 Å over 552 Cα atoms and 7.39 Å over 536 Cα atoms, respectively (Figure 3cd). The comparatively high RMSD values are likely inflated by localized structural divergence arising from low sequence identity between the proteins, rather than reflecting differences in the conserved fold (47, 48).

Together, these comparisons show that the asymmetric dimer is a conserved structural building block shared among MakA, AhlB, SmhB, XaxA/XaxB, and YaxA/YaxB. While these proteins differ in sequence, subunit composition, and oligomeric organization, they all employ a remarkably similar asymmetric dimeric architecture that serves as the foundation for their higher-order assemblies. It is worth noting that the soluble forms of XaxA/XaxB and YaxA/YaxB are predominantly α-helical and do not exhibit the β-tongue structure observed in the soluble forms of MakA, AhlB, or SmhB. Consequently, the transition from soluble XaxA/XaxB or YaxA/YaxB to the pore state requires substantially less structural rearrangement (4, 5). For YaxA, the RMSD between the soluble (PDB: 6EK7) and pore forms is 1.1 Å across 259 Cα pairs. For YaxB, the ∼60 residues in the helical head region are poorly resolved in the soluble structure (PDB: 6EK8) but are already spatially constrained by the rest of the fold, such that membrane insertion is expected to require minimal reordering of these residues into an α-helical transmembrane region, rather than large-scale rotational or translational rearrangements (5). This observation suggests that membrane insertion can impose a stronger structural constraint than the soluble state, driving convergence toward a shared pore-competent architecture across divergent toxin families.

### Computational analysis of possible transmembrane conformations of MakB and MakE

Because MakA is known to adopt both soluble and detergent-associated conformations, we investigated whether MakB and MakE possess similar membrane-insertion propensity. Kyte and Doolittle hydropathy was used to analyze and compare the hydrophobicity of the MakA, MakB and MakE sequences (38) (Figure 4a-c). As expected, MakA exhibited two strongly hydrophobic regions (hydropathy score > 2) spanning residues 196-214 and 224-237, corresponding to the β-tongue in the soluble form and the transmembrane region in both chain A and chain B of the detergent-associated form (Figure 2h, Figure S1). In contrast, MakB displayed the lowest hydrophobicity, with only residues 196-197 and 210-214 reaching hydropathy scores greater than 1.5. MakE showed intermediate hydrophobicity, with residues 201-209 and 214-223 exhibiting hydropathy scores above 1.5. A similar distribution of hydrophobicity was observed in the tripartite α-PFTs from *Bacillus cereus* (NheA, NheB, and NheC) and *A. hydrophila* (AhlA, AhlB, and AhlC), in which NheB and AhlB are analogous to MakA (strong hydrophobicity), NheC and AhlC to MakE (intermediate hydrophobicity), and NheA and AhlA to MakB (weak hydrophobicity) (6, 49, 50). Notably, AhlB is the only Ahl component with a determined pore structure, composed of asymmetric AhlB homodimers. The conservation of this hydrophobicity hierarchy across the Mak, Ahl, and The systems suggests that functional specialization among the three toxin components is also conserved.

**Figure 4.**
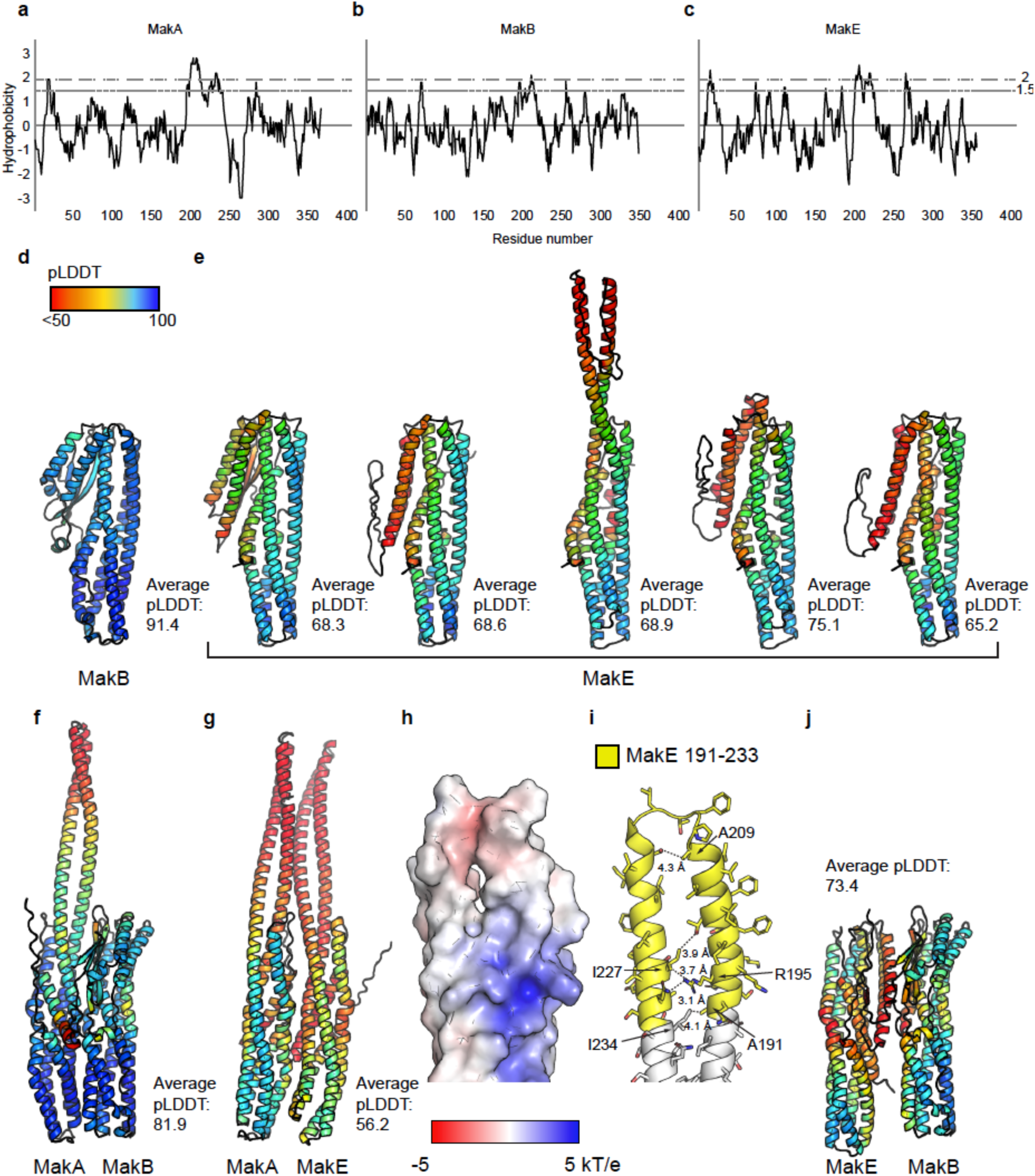
Membrane-association potential of MakA, MakB, and MakE. **a-c)** Kyte-Doolittle hydrophobicity profiles of MakA, MakB, and MakE. **d)** AlphaFold2 model of the MakB monomer generated using detergent-associated MakA as the template. **e)** AlphaFold2 model of the MakE monomer generated using detergent-associated MakA as the template. **f)** AlphaFold2 model of the MakA/MakB heterodimer generated using the detergent-associated MakA asymmetric homodimer as the template. **g)** AlphaFold2 model of the MakA/MakE heterodimer generated using the detergent-associated MakA asymmetric homodimer as the template. **h)** Surface electrostatic potential of MakE in the AlphaFold2 model of the MakA/MakE heterodimer. **i)** Potential interactions that could stabilize the transmembrane helix of MakE in the AlphaFold2 model of the MakA/MakE heterodimer. **j)** AlphaFold2 model of the MakB/MakE heterodimer generated using the detergent-associated MakA asymmetric homodimer as the template.

**Figure 5.**
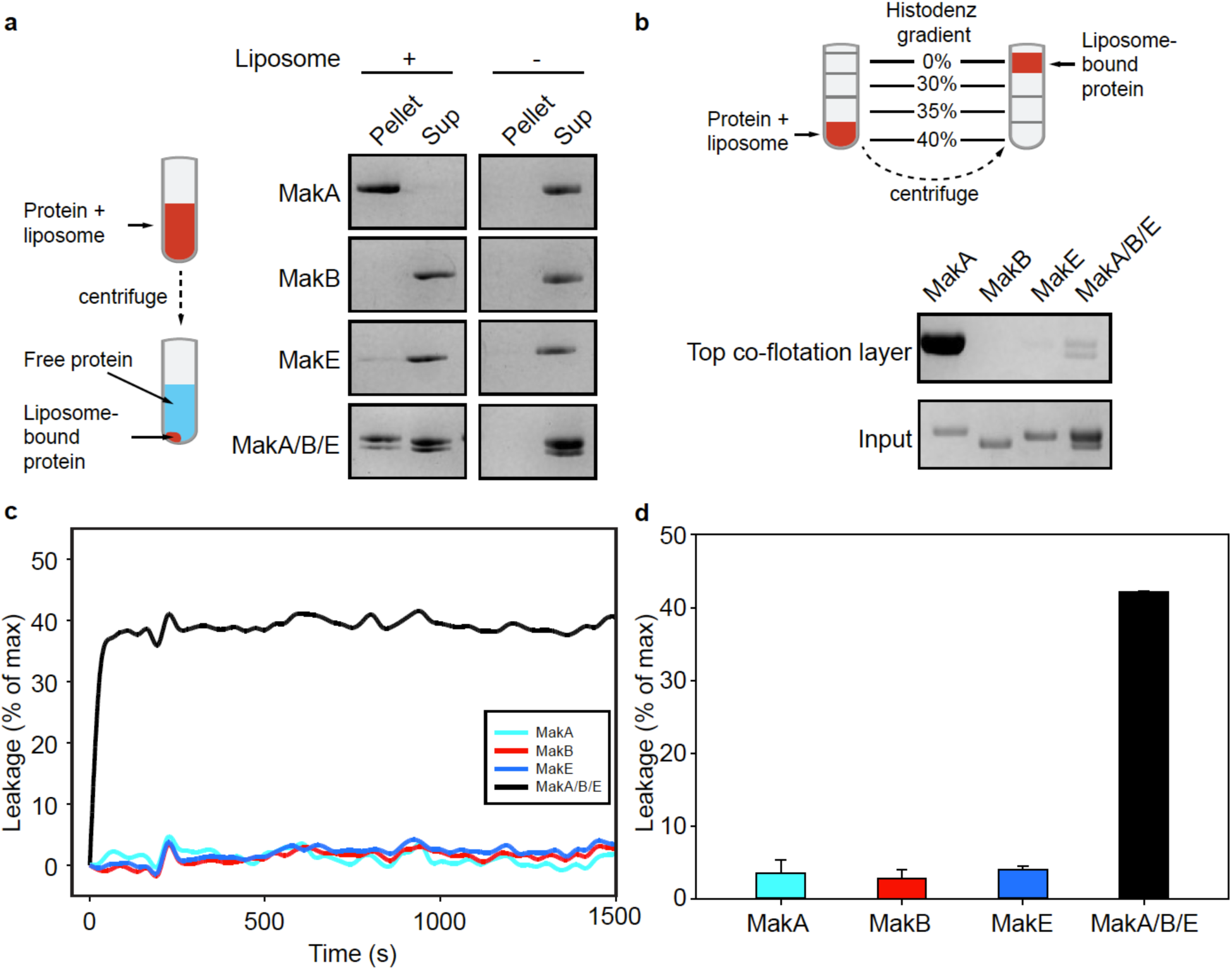
Functional characterization of MakA, MakB, and MakE using liposome assays. **a)** Liposome co-sedimentation assay to assess the membrane association of MakA, MakB, MakE, and the MakA/MakB/MakE mixture. Liposome-bound proteins are present in the pellet fraction. The right panel shows a negative control without liposomes. **b)** Liposome co-floatation assay to assess the membrane association of MakA, MakB, MakE, and the MakA/MakB/MakE mixture. Liposomes float to the top layer and their associated proteins are examined by SDS-PAGE. **c)** Liposome leakage induced by MakA, MakB, MakE, or the MakA/MakB/MakE mixture. Leakage is shown as a percentage of the maximum fluorescence signal as a function of time. Maximum fluorescence intensity was measured upon full SRB molecules release after β-OG-mediated liposome disruption. **d)** Quantification of the leakage at 1500 s. Bars represent average of the normalized fluorescence intensities performed in triplicates. Error bars represent standard deviations.

To further investigate potential transmembrane conformations of MakB and MakE, we generated AlphaFold2 models of these proteins using the detergent-associated MakA structure as a template. We also evaluated possible asymmetric homo- and heterodimeric assemblies of MakA, MakB, and MakE.

When MakA chain A or chain B was used independently as a template, AlphaFold2 consistently predicted MakB in its soluble conformation with high pLDDT scores (> 90) (Figure 4d, average pLDDT ∼91). In contrast, MakE adopted a predominantly α-helical conformation, although the position of the head (putative transmembrane) region varied among the predicted models with modest pLDDT scores (Figure 4e, average pLDDT ∼65). Using the MakA asymmetric homodimer as a template to model asymmetric homodimers of MakB or MakE did not yield plausible assemblies with well-defined dimerization interfaces.

The MakA asymmetric homodimer was then used as a template to model heterodimers between MakA, MakB, and MakE, to explore whether association with a detergent-associated MakA-like monomer could stabilize membrane-associated conformations of MakB or MakE. In the MakA/MakB heterodimer, MakB remained in its soluble conformation despite being modeled alongside detergent-associated MakA (Figure 4f, average pLDDT ∼82). In contrast, in the MakA/MakE heterodimer, MakE adopted a membrane-associated conformation that closely resembled chain B of the experimentally determined MakA homodimer, positioning its putative membrane-interacting region appropriately with respect to the lipid environment (Figure 4g, average pLDDT ∼56). Electrostatic surface analysis of the MakE head region revealed predominantly hydrophobic surfaces, consistent with a membrane-inserted state (Figure 4h). The model also suggests several plausible interhelical interactions that could stabilize the packing of the two transmembrane helices in MakE (Figure 4i). In the MakB/MakE heterodimer, although MakE adopted a predominantly α-helical conformation, its head region failed to extend into a position compatible with membrane insertion but is instead positioned between the core of MakE and the β-tongue of soluble MakB (Figure 4j, average pLDDT ∼73).

Despite low to modest pLDDT scores, the AlphaFold2 models presented here correlate well with the hydropathy analysis. Although these models contain regions of uncertain prediction, including potentially inaccurate loop conformations, they are not used to infer specific side-chain interactions or atomic-level structural details. Instead, they provide a qualitative framework to illustrate potential transition between soluble globular and membrane-associated states suggested by the hydropathy profiles. These results suggest that MakE possesses intrinsic membrane-binding features and may adopt a membrane-associated conformation alone or in the presence of an interacting partner such as a membrane-associated MakA monomer, whereas MakB appears to lack these features and is therefore expected to remain in a soluble conformation. Under this model, MakA is the principal membrane-inserting component, MakE retains the capacity to adopt a membrane-associated conformation by forming a heterodimer with MakA, and MakB contributes primarily through interactions with the tail region of MakA/MakE rather than direct membrane insertion.

### Liposome binding and disruption properties of MakA, MakB and MakE

To investigate the membrane-association and membrane-disrupting activities of the Mak proteins, we carried out liposome co-sedimentation, co-floatation and leakage assays using epithelial lipid vesicles.

Liposome co-sedimentation and co-flotation assays were first performed as complementary approaches to evaluate stable membrane association of Mak proteins (Figure 5ab). In both assays, MakA showed strong liposome association, MakE exhibited low but reproducible liposome association, and MakB showed little to no liposome binding. These differences are consistent with the membrane association propensity predicted by the sequence-based hydropathy analysis and AlphaFold2 models. The MakA/MakB/MakE mixture exhibited intermediate co-sedimentation and co-flotation, showing reduced membrane-association compared to MakA alone.

Importantly, SDS-PAGE analysis of the membrane-associated fractions revealed the presence of MakB together with a band corresponding to MakA and/or MakE, although the similar molecular weights of MakA and MakE prevented their unambiguous resolution. The detection of MakB in these fractions indicates that membrane-associated species are not composed solely of MakA or MakE, and supports the involvement of MakB in the membrane-inserted complex. Given that MakB is predicted to remain predominantly in a soluble conformation and lacks the extensive membrane-interacting features observed in MakA and MakE, it may function through decorating the soluble region of the assembled pore, as predicted by the AlphaFold2 model (Figure 4f). The reduced membrane association observed in the presence of all three proteins suggests that regulatory interactions may occur during toxin assembly which can modulate insertion kinetics, stability or stoichiometry of complex formation on the membrane. Further studies will be required to define the precise role of each protein.

Next, liposome leakage assays were performed to assess membrane permeabilization by the Mak proteins. MakA, MakB, and MakE individually exhibited little to no leakage activity (Figure 5cd). In contrast, the MakA/MakB/MakE mixture produced a strong response despite exhibiting only modest membrane association, indicating that extensive stable membrane association is not required for membrane permeabilization in the conditions tested here (Figure 5cd, Figure S5). Conversely, although MakA alone were capable of strong membrane association, it failed to induce substantial leakage, demonstrating that membrane binding alone is insufficient to disrupt the bilayer. Together, these results suggest that interactions among the Mak proteins alters their membrane interaction, leading to the formation of a functional membrane-disrupting assembly (e.g., a pore). This interpretation is consistent with the previous observation of a pore-like structure formed in liposomes prepared from *E. coli* total lipid extract following the addition of MakA, MakB, and MakE (13). The leakage could arise either from transient membrane interactions or from a relatively small population of stably membrane-associated complexes that is sufficient to induce permeabilization. The composition and architecture of the active membrane-disrupting species remain to be determined.

These observations are partially consistent with previously reported cell-based studies. In HeLa and CHO-K1 cells, MakA alone or MakA/MakE, but not MakA/MakB/MakE, were able to induce moderate cytotoxicity, whereas in Caco-2 cells and J774 cells, MakA/MakB/MakE exhibits greater toxicity than MakA, MakB or MakE, MakA/MakE or MakB/MakE (12, 13). Among these cell lines, Caco-2 is an intestinal epithelial cell line with a lipid composition most similar to that of the liposomes used in this study (51), supporting the physiological relevance of the liposome assays. Notably, J774 macrophages exhibited similar responses to Caco-2 cells despite being a distinct, non-epithelial cell type, suggesting that shared membrane properties, rather than cell type alone, may underlie the observed activity of the Mak proteins. The differences observed in HeLa and CHO-K1 cells may further reflect the contribution of cell-specific membrane composition or receptor-mediated effects (2, 15, 52, 53).

### pH-dependent activity of MakA

In this study, leakage was observed at pH 5 where MakA shows the highest membrane-association (16). This is also the same pH at which MakA induces lysosomal membrane tubulation (16). In addition, the pH-dependent membrane association of MakA may represent a defense mechanism that evolved to facilitate escape from the acidic food vacuoles of predatory protozoa (11). Although previous studies have shown that pores formed by MakA alone or by the MakA/MakB/MakE mixture can also exist at neutral pH, the link between pH and function is clear (13). Considering that protonation of histidine, aspartate, or glutamate residues at low pH is a common mechanism contributing to pH sensing in proteins, we examined all negative charged residues throughout MakA (54, 55).

The average pKa of aspartate is usually below 4, so it is unlikely to be protonated under the conditions tested previously. This leaves histidine (average pKa ∼6) and glutamate residues (average pKa ∼4-5) as potential contributors to pH-dependent responses. There are only three histidine residues in MakA, H30, H95 and H151. A recent study suggests that H30 (calculated pKa = 5.03) plays a role in pH sensing, as CD spectroscopy showed that wild-type MakA exhibits increased α-helical content under acidic conditions, consistent with a soluble β-tongue-to-transmembrane helix transition observed in this study, and the H30K mutation reduces the extent of this conformational change (56). Inspection of the MakA soluble form structure shows that H30 forms contacts with I356 and does not appear to interact with other residues. However, in the detergent-associated structure, H30 in chain B interacts with T292 in chain A and forms a hydrogen bond with S296 in chain A, suggesting that this histidine may contribute to stabilizing the asymmetric homodimer (Figure S4a). Protonation of H30 could potentially strength the hydrogen bond, assuming the Nε2 of the histidine side chain acts as the proton donor, thereby promoting the dimerization between chain A and chain B. H151 (calculated pKa = 4.64) exhibits a similar interaction pattern to H30. In the soluble form, H151 interacts with S281 and S282 within the same chain. In the detergent-associated structure, H151 in chain A forms a hydrogen bond with N34 in chain B, which could be strengthened at lower pH (Figure S4a). However, the H151K mutant was not tested in the study due to low protein stability. H95 (calculated pKa = 5.94) interacts with residues within the same chain in both the soluble and detergent-associated forms, and the H95K mutant behaves similarly to the wild-type protein, making it unlikely to be directly involved in pH sensing. It should be noted that strengthening a single hydrogen bond may not be sufficient to explain the functional effect. In addition, the results observed with the H30K mutation may simply reflect the longer side chain of lysine introducing steric clashes at the dimerization interface and disrupting efficient packing required to stabilize the asymmetric MakA homodimer. In the absence of this stabilizing interaction, MakA may remain predominantly in the soluble form with lower α-helical content.

Further inspection of the soluble form of MakA identified two glutamate residues of interest, E357 and E187. E357 (calculated pKa = 4.07), located in the last β-strand of the β-tongue structure, can form a hydrogen bond with side chain of W243 in the post-β-tongue helix (Figure S6a). E187 (calculated pKa = 4.87), located at the pre-β-tongue helix, can form a hydrogen bond with S286, which lies within the core soluble domain of MakA (Figure S6b). Both interactions likely contribute to “locking” the β-tongue in the soluble conformation. Protonation of glutamate residues would weaken the hydrogen bond, thus destabilizing the lock and facilitating the transition from soluble to the transmembrane conformation.

It is important to point out that protonation of histidine and glutamate residues not only affects a subset of hydrogen bonds, but can also disrupt stronger electrostatic interactions such as salt bridges between E173^chain A^/R183^chain B^ and K329^chain A^/E96^chain B^ at the dimer interface, and can alter the overall electrostatic potential. Therefore, a more carefully designed mutagenesis study should be carried out to identify the key pH-sensing residues in MakA. It is also possible that no single residue acts as a dominant pH sensor, but instead a distributed network of small changes in hydrogen bonding and electrostatic interactions collectively lowers the energetic barrier for the transition to the transmembrane conformation.

### Evolutionary analysis of bacterial pore proteins

MakA shares intermediate to low sequence similarity with other α-PFTs (<40% sequence similarity). However, the MakA asymmetric homodimer exhibits strong structural similarity to both the homodimeric pore-forming building blocks of AhlB and SmhB, and the heterodimeric pore-forming building blocks of XaxAB and YaxAB. This combination of low sequence conservation and high structural similarity suggests a shared ancestral origin followed by early divergence. To further examine the relationships among these toxins, we performed a phylogenetic analysis including toxins with sequence and/or structural similarity. For genes involved in host-pathogen interactions, horizontal gene transfer is frequent; consequently, the phylogenetic tree presented here primarily reflects the evolution of the toxins themselves rather than that of the organisms harboring them.

The phylogenetic tree shows that MakA (Figure 6, green) and MakE (Figure 6, magenta) cluster as sister paralogs, with MakB (Figure 6, cyan) representing a long, possibly more basal branch. This topology is consistent with an initial duplication separating MakB from the MakA/MakE ancestor, followed by a second duplication giving rise to MakA and MakE. YaxA/XaxA and YaxB/XaxB also form well-supported orthologous clades, consistent with a single ancestral duplication producing the A- and B-type subunit lineages prior to the divergence of toxins in *Yersinia* and *Xenorhabdus*. Homologs from additional bacterial toxins within each clade further indicate that these duplication events predate the diversification or horizontal transfer of such genes (57). Because the Mak proteins branch separately from the YaxA/XaxA and YaxB/YaxB lineages, the Mak system differs fundamentally from these bipartite systems in both composition and evolutionary history. Although MakA and MakE together are sister to the XaxB/YaxB clade, this relationship does not indicate direct one-to-one orthologous relationship between MakA/MakE and the B-type toxin subfamilies.

**Figure 6.**
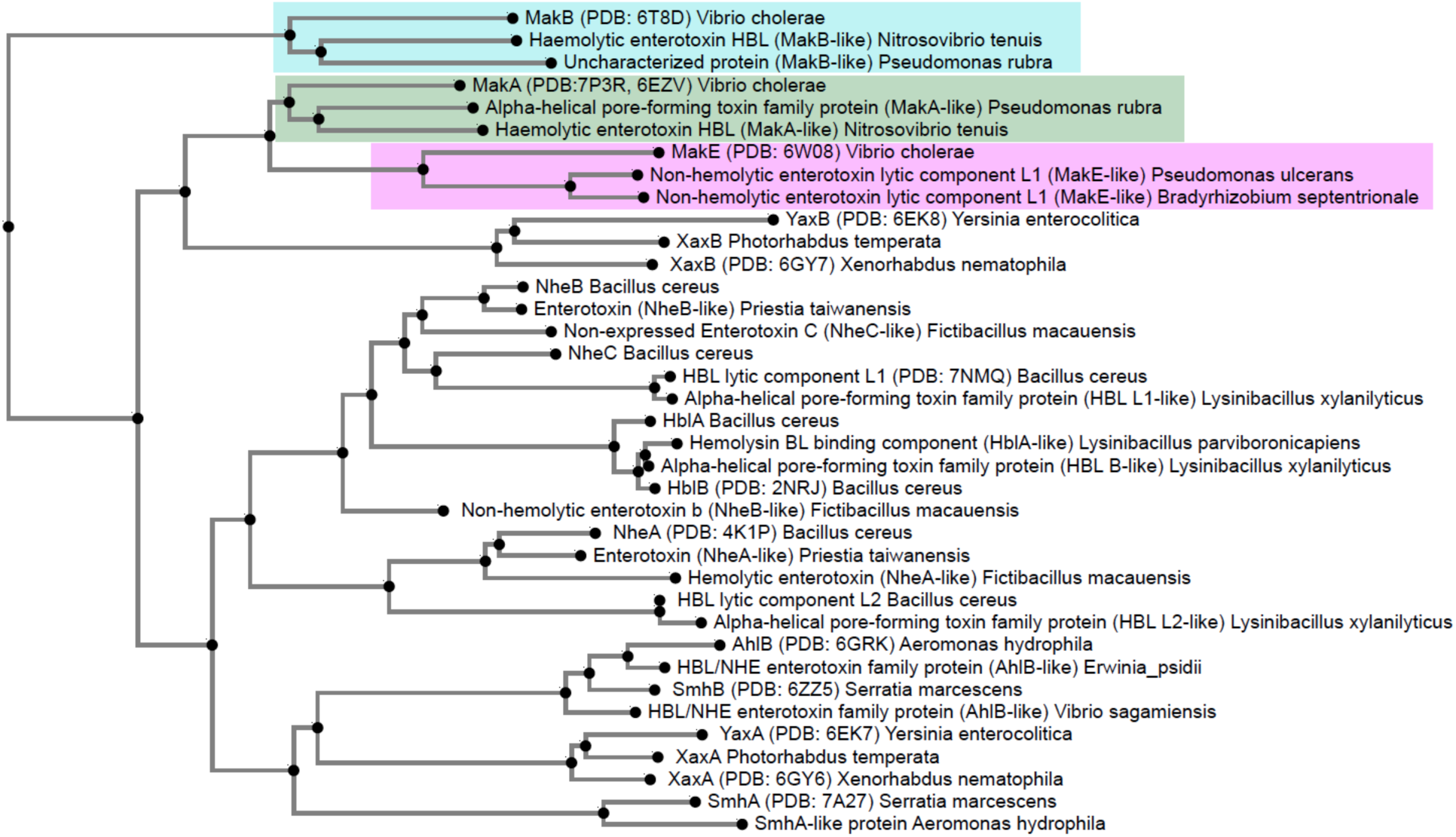
Phylogenetic tree of selected α-pore-forming toxins from various species. MakA (green) and MakE (magenta) cluster as sister paralogs, whereas MakB (cyan) forms a more basal branch, consistent with sequential gene duplication. MakA, MakB, and MakE form a distinct lineage within the α-pore-forming toxin superfamily and do not show a one-to-one orthologous relationship with other toxin families.

Notably, despite being a tripartite system, Mak differs from other three-component pore-forming toxins in its evolutionary origin and subunit organization as well. For example, the Nhe system from *B. cereus* (NheA, NheB, NheC) falls into separate phylogenetic lineages from the Mak proteins, indicating that tripartite architecture arose independently in these two systems through distinct duplication trajectories (58).

Together, these observations suggest that multicomponent pore-forming toxins do not conform to a single conserved evolutionary pattern of subunit architecture, but instead have repeatedly emerged through independent gene duplication events followed by functional divergence.

## Discussion and conclusion

In this study, we determined a detergent-associated structure of MakA that assembles as a trimer of asymmetric homodimers. Notably, the same asymmetric homodimer is also present in the previously reported helical form of MakA despite the direrent higher-order organization of the two assemblies (16). The recurrence of this asymmetric homodimer in independent MakA structures suggests that it represents a biologically relevant assembly rather than an artifact of detergent stabilization. Consistent with this interpretation, the MakA homodimer displays significant structural similarity to the heterodimeric pore-building units of the distantly related homologs XaxA/XaxB and YaxA/YaxB.

Sequence analysis, hydropathy profiling, AlphaFold2 modeling, and liposome assays further support distinct roles for the three Mak proteins during membrane interaction. MakA contains the most pronounced hydrophobic region and readily adopts a transmembrane conformation. MakE possesses an intermediate hydrophobic patch and is capable of inserting into liposomes, despite its low membrane association. In contrast, MakB exhibits substantially lower hydrophobicity and is consistently predicted to remain in its soluble conformation, suggesting that it is less likely to contribute directly to the transmembrane portion of the complex.

The strong hydrophobic character of MakA makes it a plausible initiator of membrane insertion, prompting an update to the model described by Herrera *et al* (12). Thus, we propose that the MakA trimer-of-dimers represents an early intermediate in pore assembly, or a pre-pore state. The tripod-like architecture provides ample space for the recruitment of additional protein components, while the relatively small, buried surface area between asymmetric dimers stabilizes the assembly yet remains permissive to the subsequent incorporation and rearrangement of additional dimers during pore formation.

Like YaxA in the YaxAB system, the pre-pore MakA assembly may also facilitate the ordered incorporation of additional subunits. In YaxAB, YaxA is readily inserted into membranes, whereas YaxB requires pre-existing membrane-associated YaxA for insertion, indicating that the sequence of subunit incorporation can critically influence the final pore architecture (5). Such a mechanism could prevent premature or nonspecific membrane engagement and promote assembly only under favorable conditions, such as upon reaching a suricient local concentration or encountering an appropriate membrane environment.

It is also worth mentioning that uncontrolled formation of the pre-pore MakA may be unfavorable for efficient pore formation, as excessive nucleation events could compete with productive pore assembly; therefore, their formation likely needs to be regulated. Previous studies demonstrated that MakB alone can reduce MakA-mediated cytotoxicity in cultured cells (12, 13). Consistent with these observations, our liposome co-sedimentation and co-floatation assays showed reduced MakA membrane association when MakB was present. These findings raise the possibility that MakB functions as a regulatory component that interacts with soluble MakA and/or MakE to modulate membrane insertion kinetics for proper pore formation. This hypothesis is supported by a genetic study of *V. cholerae* survival in *Tetrahymena pyriformis* using deletion and complementation of *makA*, *makB* and *makE*. In this study, both *makA* and *makE* are required for *V. cholerae* survival within the protozoan food vacuoles, likely by inserting into the phagosomal membrane and disrupting vacuolar function, thereby promoting resistance to digestion. In contrast, all three proteins are required to kill *T. pyriformis*, consistent with a model in which MakB is required for assembly of mature pores that enable *V. cholerae* to escape into the cytoplasm, ultimately leading to host cell death (11).

Guided by the architectures of the AhlB and XaxAB pores, we generated a hypothetical model of the Mak pore by replacing their pore-building units with the experimentally determined MakA homodimer (Figure 7a-c). Based on this model, we further speculate that membrane insertion of additional assembly units into the pre-pore creates new intermolecular interactions between neighboring subunits, which in turn drive structural rearrangements that promote pore formation (Figure 7d-f). For example, increased contacts within both the membrane-inserted and soluble regions could induce rotational and translational movements of the building blocks, gradually reorienting them toward the central axis and reducing the angle between each building block and the pore axis, ultimately generating an open transmembrane channel.

**Figure 7.**
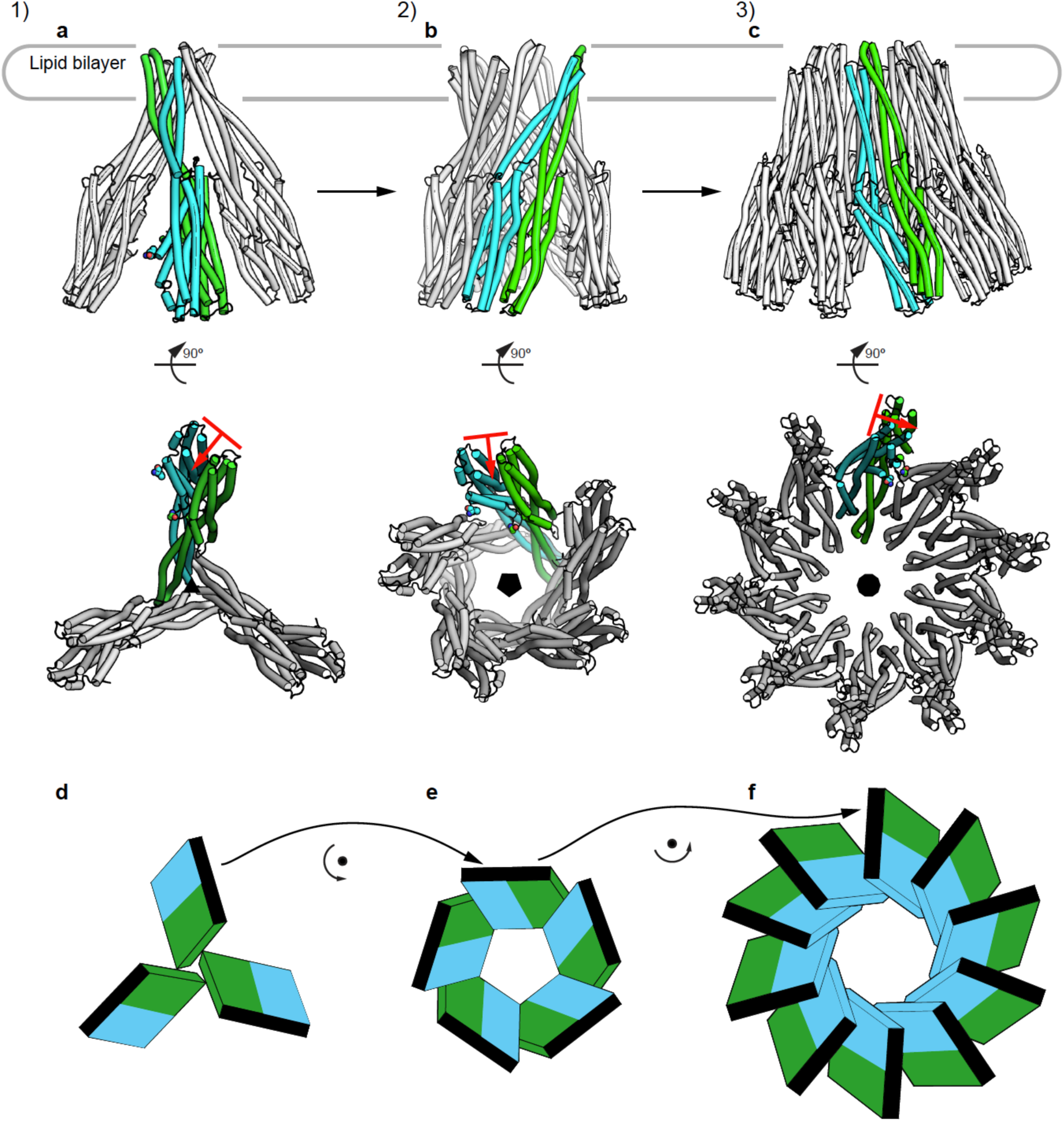
Model of MakA pore formation. **a-c)** Proposed pore assembly initiated from a trimer of dimers. The orientation of each asymmetric dimer building block relative to the symmetry axis changes as additional subunits are incorporated (indicated by the red line and arrow). **d-f)** Illustration of subunit rotation during pore growth. Following membrane insertion of MakA and/or MakE into the initial trimer of dimers, intermolecular interactions between neighboring assembly units drive progressive rotational and translational rearrangements that reorient subunits toward the pore axis, decreasing the angle between each building block and the central channel and promoting formation of an open transmembrane pore. In this model, MakB may remain associated with the soluble region or function outside the mature pore complex.

This model should be regarded as a working framework rather than a definitive structural description. The mature Mak pore may exhibit direrent symmetries, alternating MakA, MakB, and MakE subunits in the assembly, or variable oligomeric stoichiometries, as observed for the XaxAB and YaxAB pores, which have been reported to contain 13-15 and 8-12 subunits, respectively.

To better understand how the Mak pore assembles and functions, additional experimental data will be required to test the mechanistic model proposed here. In particular, structural approaches in a native-like lipid environment will be important to confirm whether the asymmetric dimer arrangement observed in detergent is retained upon membrane association, and whether additional conformational changes occur during lipid insertion. Membrane curvature and a broader range of lipid compositions could also be tested to identify possible signals that triggers the Mak pore formation, including membrane models that mimic protozoan food vacuoles, given Mak proteins’ proposed role in promoting bacterial survival against predatory protozoan engulfment (11, 15, 59, 60). Such studies could be complemented by time-resolved assembly assays in which MakA, MakB, and MakE are sequentially added to lipid vesicles in different orders to dissect the dependence of membrane insertion and assembly dynamics (61). Targeted mutagenesis of residues in the putative transmembrane and inter-dimer interfaces of MakA, MakB, and MakE would further clarify their roles in membrane insertion, stabilization of the asymmetric dimer architecture, and subunit-specific interactions. The mutagenesis results should also be interpreted with caution because the same MakA sequence exists in three different structures, and MakB and MakE may likewise adopt multiple conformations.

Although the architecture and mechanism of mature pore assembly remain to be determined, our structure supports the asymmetric MakA dimer as a candidate building block of the pore. Pharmacological intervention targeting this building block, either by disrupting the dimer interface or by preventing its formation from soluble MakA, could interfere with membrane insertion and halt pore assembly at an early stage.

The MakA assembly described here also highlights a limitation of current structure prediction methods. Although AlphaFold2 accurately predicts the soluble structures of MakA, MakB, and MakE, it does not readily predict the experimentally observed MakA asymmetric homodimer. The detergent-associated MakA assembly involves two copies of the same protein adopting distinct conformations within a single complex, which is a scenario that remains challenging for current prediction algorithms. Similar difficulties have been reported for other dynamic protein assemblies that undergo extensive conformational rearrangements (62). These observations emphasize the continued importance of experimental structural determination for characterizing alternative conformational states and higher-order assemblies that are not readily captured by sequence-based prediction methods.

## Acknowledgement

The research reported in this publication was supported by the National Institute of Allergy and Infectious Diseases (NIAID) of the National Institutes of Health (NIH), Department of Health and Human Services, under contracts HHSN272201200026C (K.J.F.S., A.J., Z.O.) and 75N93022C00035 (K.J.F.S., A.J., D.B.). Additional support was provided by NIAID under award numbers K99AI167819 (A.H) and R37AI092825 (K.J.F.S.); the National Institute of General Medical Sciences (NIGMS) of NIH under award numbers R44GM137671 (R.B., Z.O.) and R35GM145365 (Z.O.); and the National Institute of Neurological Disorders and Stroke (NINDS) of NIH under award number R35NS097333 (J.R.). The research reported in this publication was financed 100% with federal funds, totalling an estimated $750,000, and 0% with nongovernmental funds ($0).

The content is solely the responsibility of the authors and does not necessarily represent the original views of the National Institutes of Health. This manuscript is the result of funding in whole or in part by the National Institutes of Health (NIH). It is subject to the NIH Public Access Policy. Through acceptance of this federal funding, NIH has been given a right to make this manuscript publicly available in PubMed Central upon the Original Date of Publication, as defined by NIH.

We thank the Cryo-Electron Microscopy Facility (CEMF) at UT Southwestern Medical Center for maintaining the electron microscopes. The facility is supported by grants RP170644 and RP220582 from the Cancer Prevention & Research Institute of Texas (CPRIT).

## Author contributions

Conceptualization: A.J., K.J.F.S., J.R., D.B., Z.O.; Methodology: Y.G., B.Q., K.P.S., R.B., J.R., D.B. and Z.O.; Investigation: R.J. and Y.K. performed protein expression and purification; B.Q., T.E. and D.B. prepared cryo-EM grids; B.Q. and D.B. collected cryo-EM data; Y.G., B.Q. and Z.O. performed single-particle reconstruction; Y.G. validated models and maps; Y.G. performed structure-guided analysis and AlphaFold modeling; K.P.S. performed liposome assays; Y.G., R.B. and Z.O. performed phylogenetic analysis; Y.G. and Z.O. worked on mechanistic models. Supervision: A.J., K.J.F.S., J.R., D.B., Z.O.; Writing - original draft: Y.G.; Writing - review and editing: Y.G., K.P.S., A.H., R.J., Y.K., K.J.F.S, J.R., D.B., Z.O.; Financial support: A.H., R.B., A.J., K.J.F.S., J.R., D.B., Z.O.

## Competing interests

Y.G., R.B., D.B. and Z.O. are cofounders of Ligo Analytics. Y.G. serves as the CEO of Ligo Analytics. Y.G. and R.B are currently employed by Ligo Analytics. Z.O. is a co-founder of HKL Research. K.J.F.S. has a significant financial interested in Situ Biosciences, a contract research organization that conducts research unrelated to this work.

---

**Supplementary Figure 1.**
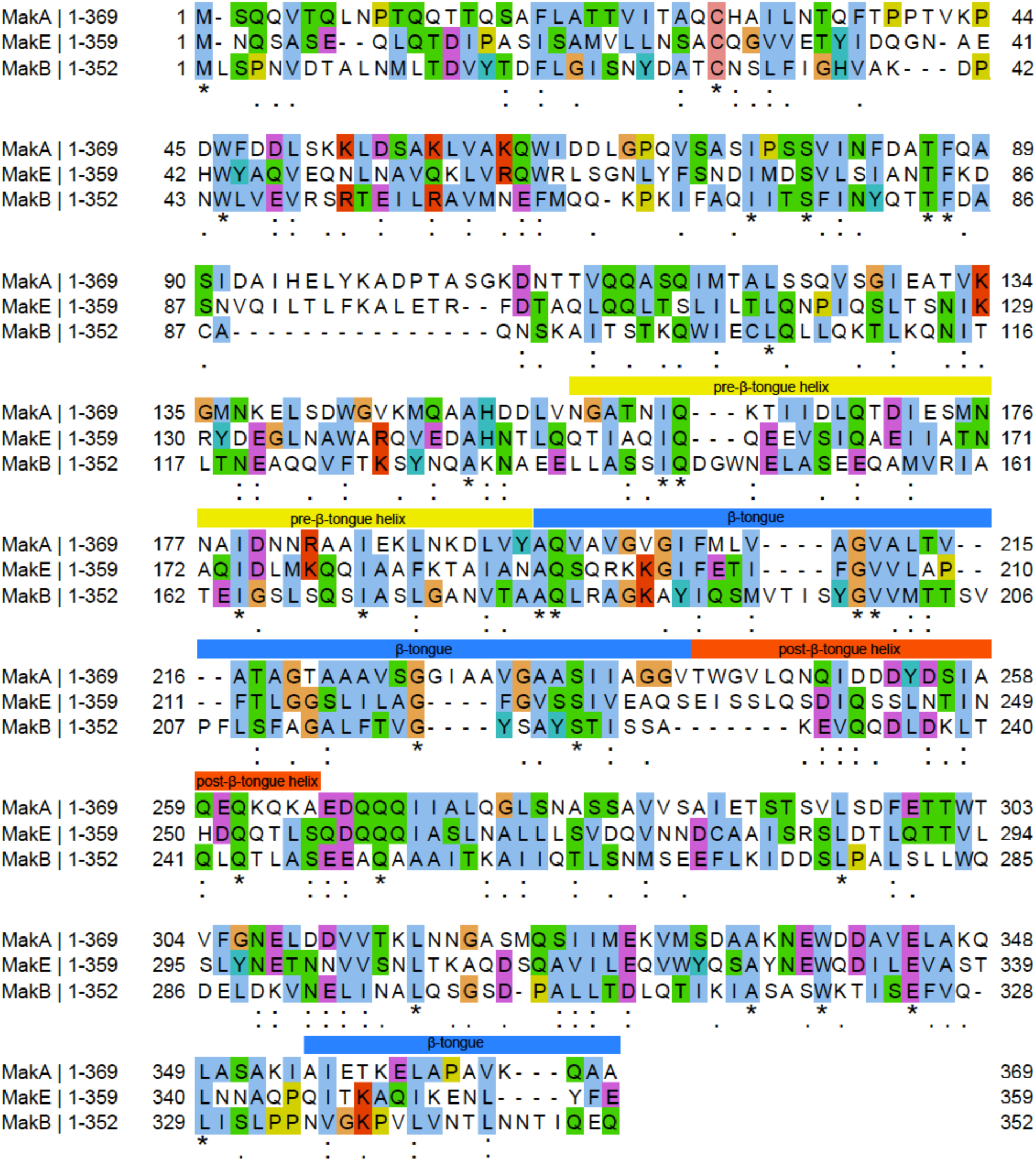
Sequence alignment of *V. cholerae* MakA, MakB and MakE. Alignment is colored by the Clustal color scheme. Conservation is independently indicated below the alignment using Clustal notation: asterisk (*) indicates fully conserved residues, colon (:) indicates strongly similar residues, and period (.) indicates weakly similar residues.

**Supplementary Figure 2.**
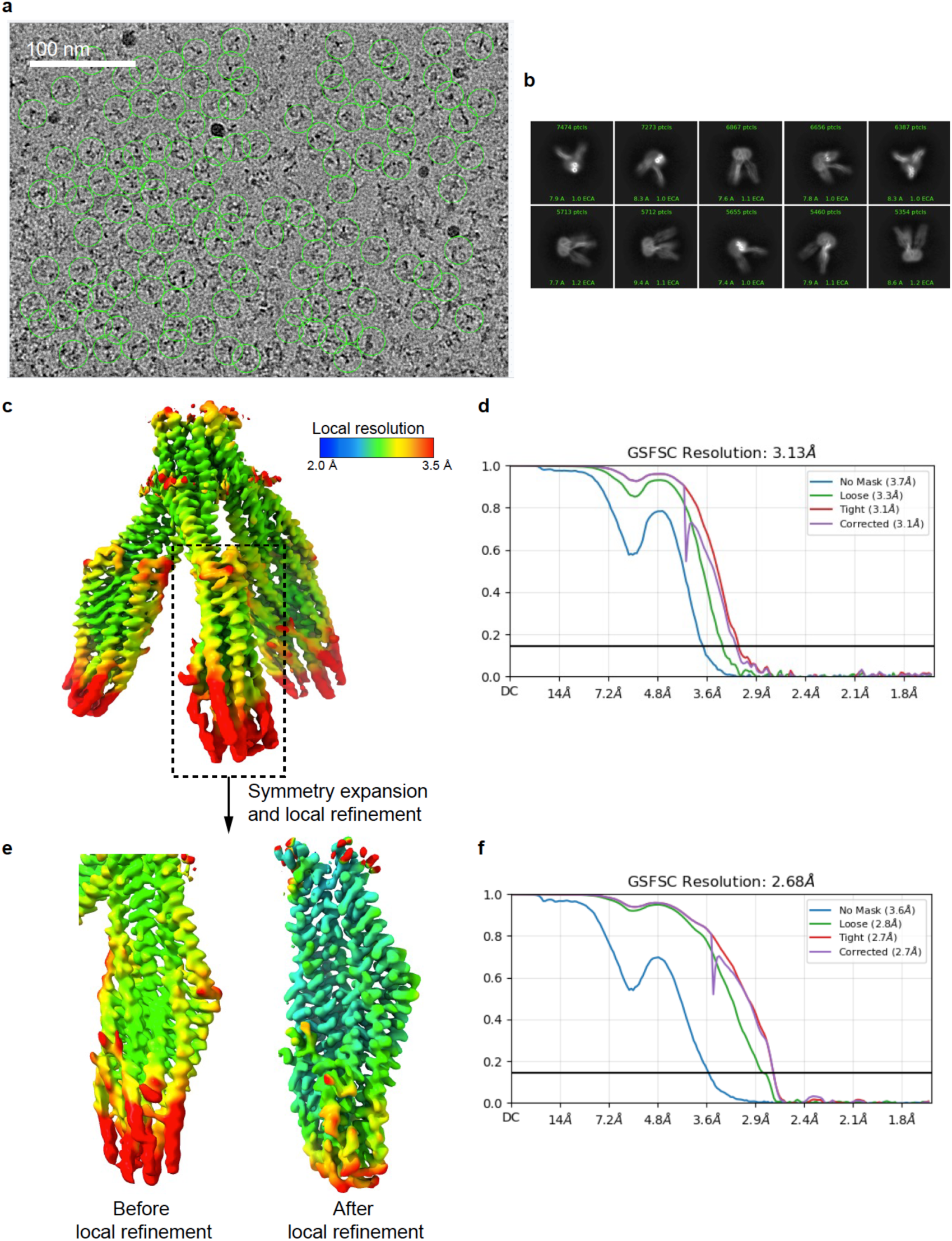
Cryo-EM workflow. **a)** Motion-corrected micrograph. Particles are highlighted with green circles. **b)** Selected 2D classes. **c)** Global reconstruction showing high local resolution in the head region and lower local resolution at the tip of the tail region due to motion. **d)** FSC curve of the global reconstruction. **e)** Side-by-side comparison of the tail region before and after local refinement using symmetry-expanded particles. **f)** FSC curve of the locally refined tail region.

**Supplementary Figure 3.**
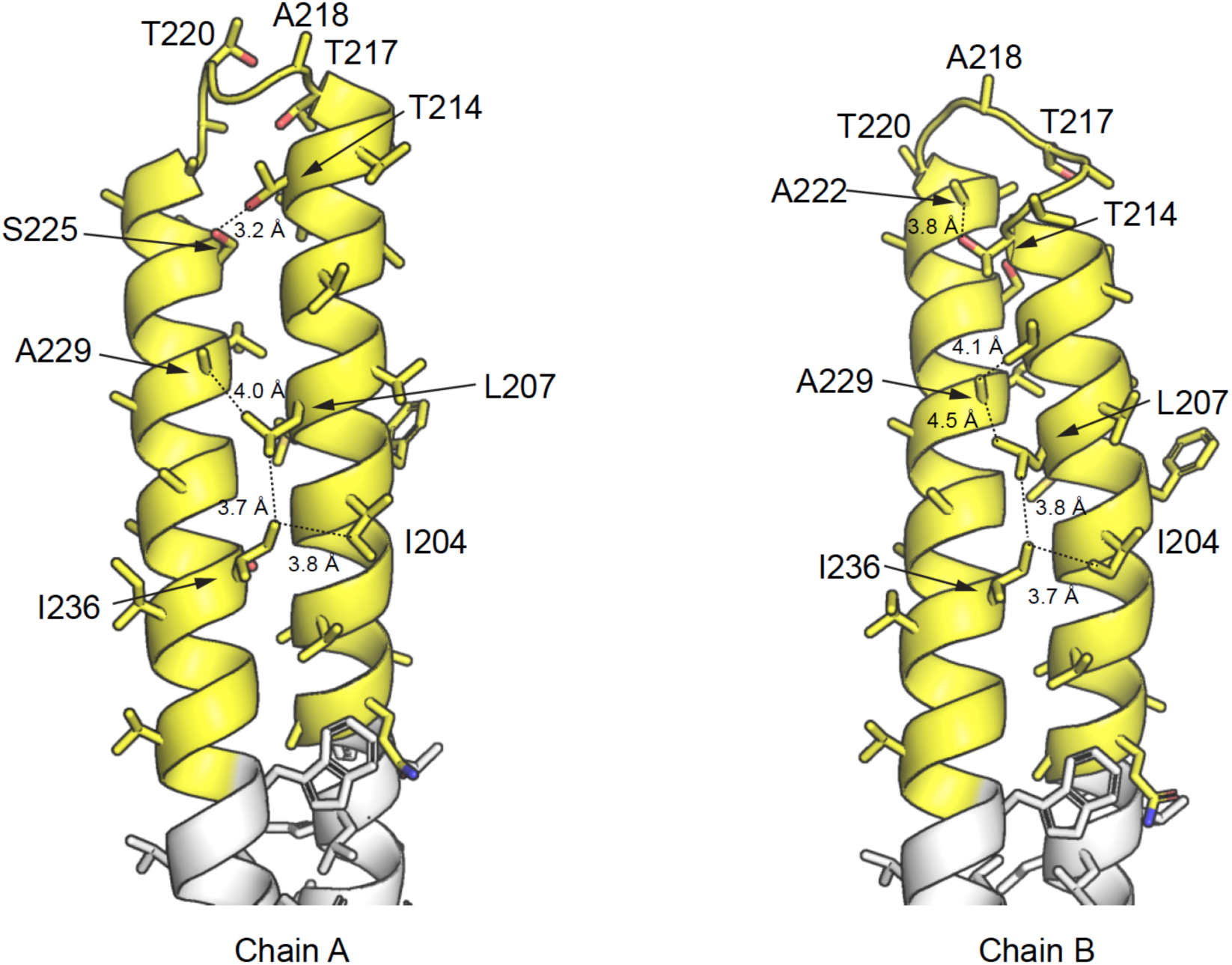
The transmembrane regions of detergent-associated MakA chains A and B adopt different conformations. Transmembrane residues are in yellow.

**Supplementary Figure 4.**
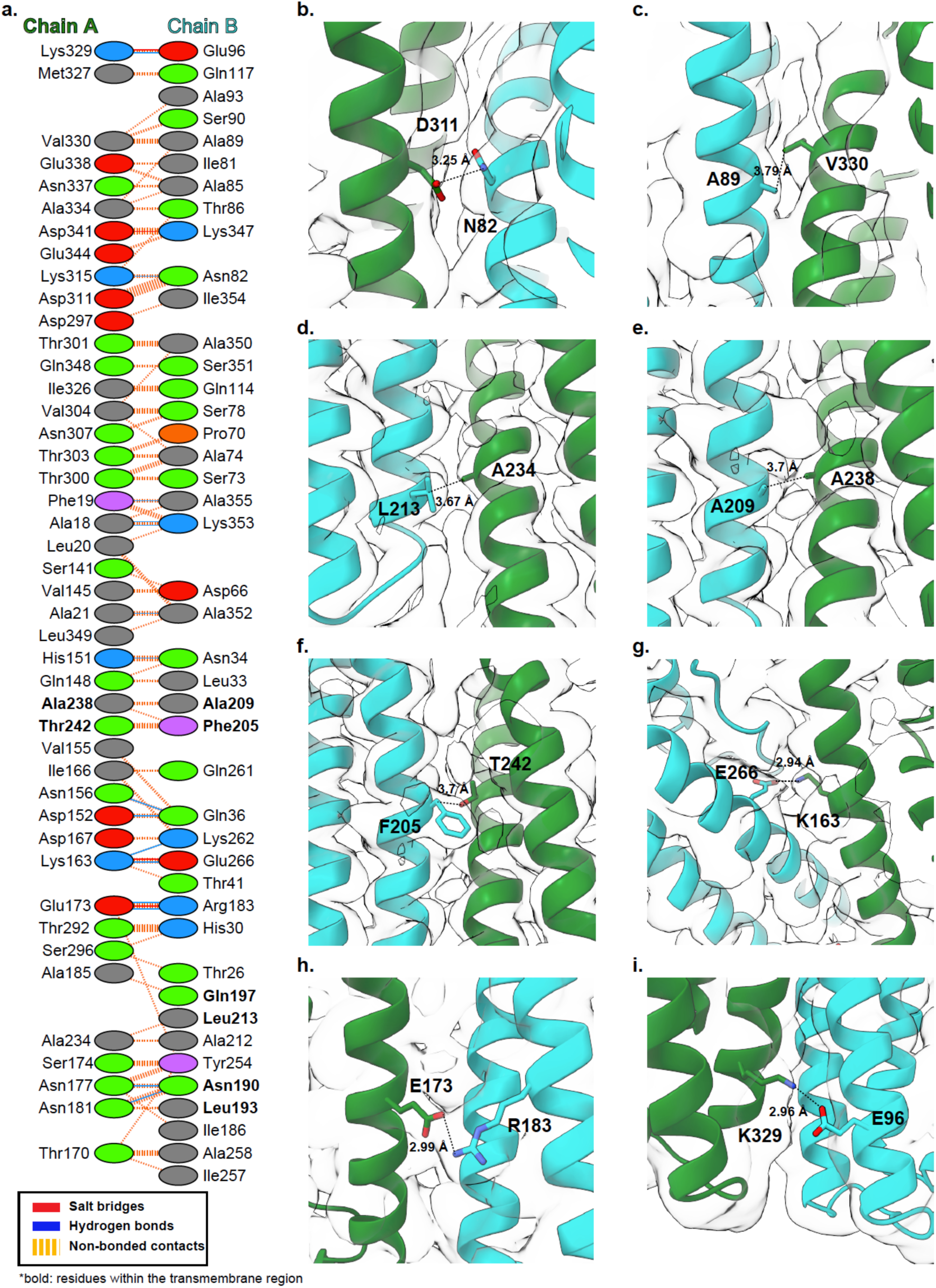
Interactions within an asymmetric dimer. **a)** Types of interactions between chain A and chain B within an asymmetric dimer. Residues in the transmembrane region are shown in bold. **b-i)** Unsharpened cryo-EM density map and fitted structural segments of the MakA asymmetric dimer.

**Supplementary Figure 5.**
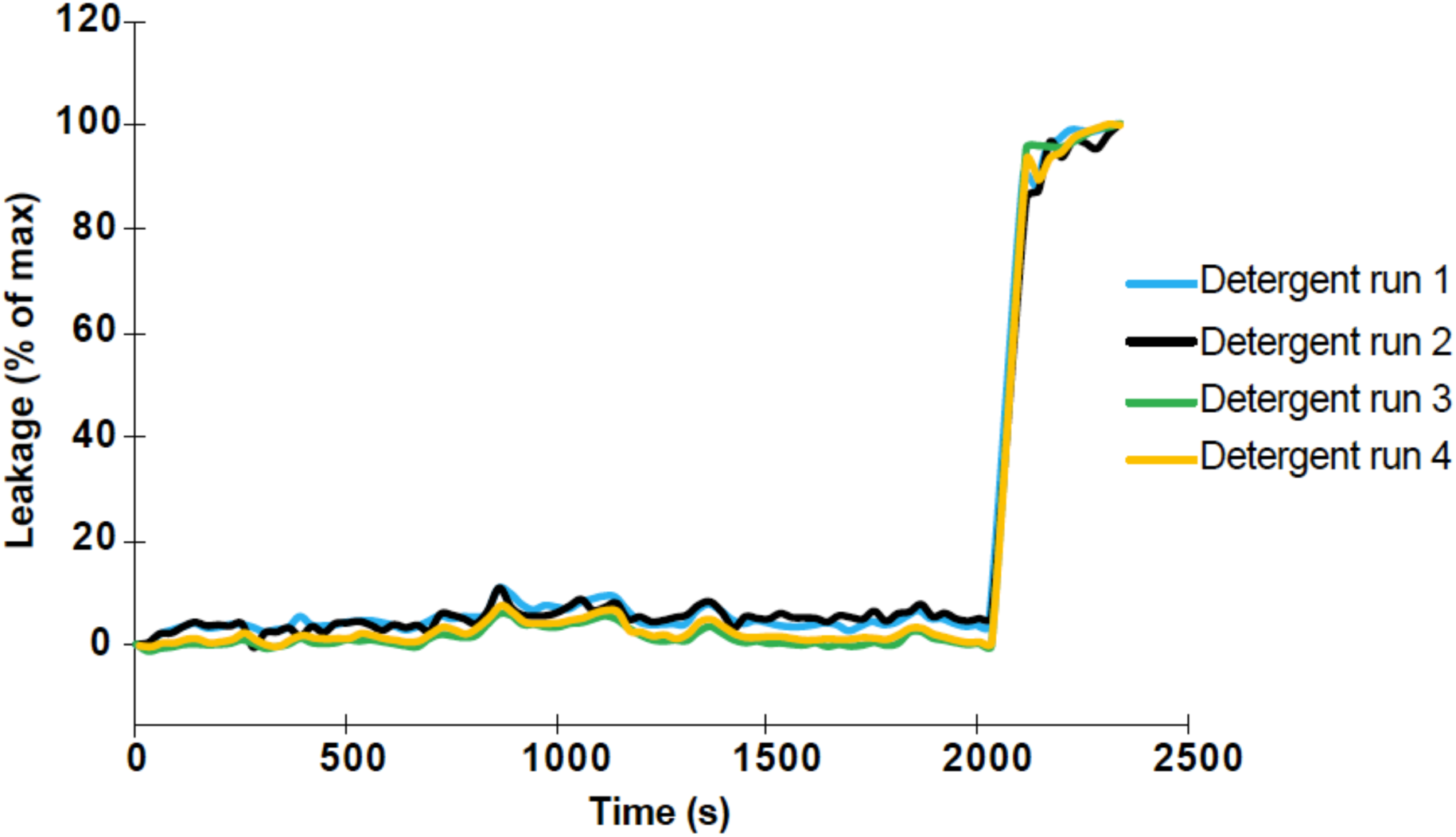
Liposome stability in the presence of detergent. Residual detergent remaining after liposome preparation did not induce detectable leakage. At 2,000 s, excess β-OG was added to disrupt the liposomes, resulting in complete SRB release and maximum fluorescence intensity.

**Supplementary Figure 6.**
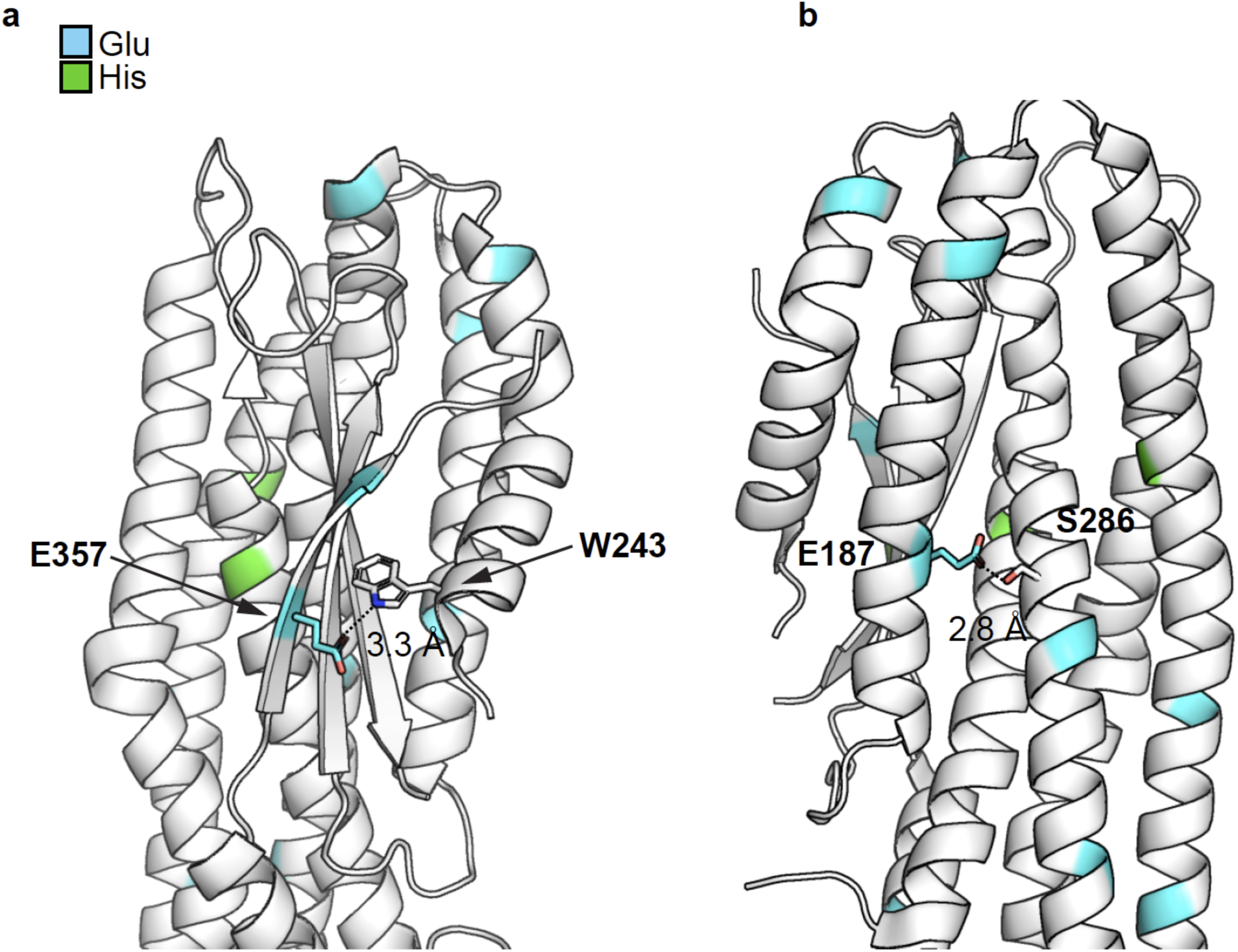
Interactions in the MakA soluble form (PDB: 6EZV) involving glutamate residues and the β-tongue. Glutamate residues are highlighted in cyan. Histidine residues are highlighted in green. Hydrogen-bond forming residues are shown as sticks.

